# Multiparameter Quantitative Analyses of Diagnostic Cells in Brain Tissues from Tuberous Sclerosis Complex

**DOI:** 10.1101/2024.02.19.581031

**Authors:** Jerome S. Arceneaux, Asa A. Brockman, Rohit Khurana, Mary-Bronwen L. Chalkley, Laura C. Geben, Matthew Vestal, Muhammad Zafar, Sarah Weatherspoon, Bret C. Mobley, Kevin C. Ess, Rebecca A. Ihrie

## Abstract

The advent of high-dimensional imaging approaches offers innovative opportunities to molecularly characterize diagnostic cells in disorders that have previously relied on histopathological definitions. One example of such disorders is tuberous sclerosis complex (TSC), a developmental disorder characterized by systemic growth of benign tumors. Within resected brain tissues from patients with TSC, detection of abnormally enlarged balloon cells (BCs) is pathognomonic for this disorder. Though BCs can be identified by an expert neuropathologist, little is known about the specificity and broad applicability of protein markers for these cells, complicating classification of proposed BCs identified in experimental models of this disorder. Here, we report the development of a customized machine-learning workflow (<u>Ba</u>lloon <u>Iden</u>tifier; BAIDEN) that was trained to prospectively identify BCs in tissue sections using a histological stain compatible with high-dimensional cytometry. This approach was coupled to a custom antibody panel and 36-parameter imaging mass cytometry (IMC) to explore the expression of multiple previously proposed BC markers and develop a descriptor of BC features conserved across multiple tissue samples from patients with TSC. These findings comprise a toolbox and dataset for understanding the abundance, structure, and signaling activity of these histopathologically abnormal cells, and an example case of how such tools can be developed and applied within human tissues.

## Introduction

The advent of high-dimensional approaches for studying tissue specimens, including spatial transcriptomics and multiparameter imaging cytometry (e.g., CODEX, MIBI, and IMC) has greatly expanded opportunities to perform detailed molecular analyses of cell populations that were previously identified using morphological features (Chang et al., 2017; Leelatian et al., 2017; Nowicka et al., 2017; Keren et al., 2019; Black et al., 2021; Brockman et al., 2021; Karimi et al., 2023). Such approaches typically focus on smaller regions of interest, but this can be a challenge when the population of interest is rare or sparsely spread across a tissue specimen. As patient-derived models also expand in scope and complexity, a second question is how best to quantitatively compare unusual cell phenotypes present in these systems to the patient tissues they are intended to model. One tissue where many such models have been developed is the brain, with a particular emphasis on developmental or genetic disorders that are studied using models such as cerebral organoids (Lancaster et al., 2013; Blair et al., 2018; Sloan et al., 2018; Eichmuller et al., 2022).

Focal cortical dysplasias (FCDs) are neurodevelopmental disorders characterized by disorganization of the cerebral cortex, including disordered lamination and the presence of multiple dysmorphic cells; these disorders have an incidence of 1 in 2000 live births annually (Neligan and Sander, 2011; Crino, 2015; Blumcke et al., 2017; Blumcke et al., 2021a; Blumcke et al., 2021b). Nearly two-thirds of individuals with FCD have intractable epilepsy, and FCDs account for about 17% of the pediatric surgical population, underscoring an urgent medical need for therapeutic targets (Thiele, 2010; Crino, 2015; Baldassari et al., 2019). Nearly all identified causative mutations for FCDs (∼90%) affect proteins interacting with or within the mammalian / mechanistic target of rapamycin (mTOR) pathway, a central regulator of cell size and growth (Lipton and Sahin, 2014; Crino, 2016; Saxton and Sabatini, 2017; Baldassari et al., 2019). Currently, FCDs associated with mutations in mTOR pathway-associated proteins other than *TSC1* (hamartin) or *TSC2* (tuberin) are categorized as non-TSC-associated FCDs; this class includes mutations in *MTOR*, *AKT*, *RHEB*, *DEPDC5*, *PTEN*, and *PIK3CA* (Baulac et al., 2015; D’Gama et al., 2017; Ribierre et al., 2018; Baldassari et al., 2019). Heterozygous mutations in either *TSC1* or *TSC2* are categorized as TSC-associated FCD and are found in patients with tuberous sclerosis complex (TSC), an autosomal dominant neurodevelopmental disorder (Camposano et al., 2009; Crino, 2013; Baldassari et al., 2019). TSC affects about 1 in 6000 live births annually and is characterized by the growth of benign tumors, also termed hamartomas, in many organs including the brain (Zimmer et al., 2020). Brain hamartomas in TSC include subependymal giant cell astrocytomas (SEGAs), which can cause increased intracranial pressure and require emergent neurosurgical intervention, and tubers, which cause significant morbidity including epilepsy and cognitive impairment (Mizuguchi and Takashima, 2001; DiMario, 2004; Zimmer et al., 2020). Patients with TSC also frequently exhibit developmental delay, intellectual impairment, autism spectrum disorder (ASD), and TSC-associated neuropsychiatric disorders (TANDs) (Barkovich et al., 2015; De Vries et al., 2015).

At the cellular level, TSC-associated FCD is distinguished by the presence of balloon cells (BCs), atypical cells identified by an enlarged, ovoid soma and eccentrically placed nucleus (Urbach et al., 2002; Lamparello et al., 2007; Crino, 2013). Though these cells are a diagnostic feature of TSC, their possible origins and properties have been relatively unexplored until the advent of mouse models and patient-derived *in vitro* models (e.g., cerebral organoids). Multiple such models have been reported to contain abnormal cells that histologically resemble BCs; however, the molecular profile of BCs in patient tissues has not been systematically described, complicating such comparisons (Li et al., 2017; Blair et al., 2018; Eichmuller et al., 2022; Karalis et al., 2022; Kovermann et al., 2022; Wu et al., 2022).

In prior studies of patient tissues, many proteins have been postulated to selectively identify BCs, including proteins associated with progenitor cells, mature neuronal and glial cells, and phosphorylation events driven by mTOR kinase activity (Hilbig et al., 1999; Mizuguchi et al., 2002; Urbach et al., 2002; Lamparello et al., 2007; Orlova et al., 2010; Yasin et al., 2010; Ruppe et al., 2014; Li et al., 2015; Rossini et al., 2017; Day et al., 2020; Gelot and Represa, 2020; Wu et al., 2022). However, these studies used single- or double-channel immunohistochemistry, limiting the ability to identify coexpression patterns, and it is not yet clear how uniformly any given protein is expressed in BCs across either individual tubers or patient cohorts (Urbach et al., 2002; Palmini et al., 2004; Najm et al., 2022).

An additional factor limiting efforts to study BCs in the numbers needed for statistically robust descriptions is the challenge of manually identifying these cells in a large, resected sample of hundreds of square millimeters, which can be a laborious and time-intensive process. This study introduces a trainable, machine-learning approach to efficiently identify BCs from whole-slide images and couples these data to analyze BCs in tuber tissue via imaging mass cytometry (IMC), allowing for the rapid identification of a sufficient number of BCs for statistical analysis across large sections. A custom antibody panel was developed in parallel to simultaneously quantify 30+ protein species of interest in patient tissue sections. Using this approach across samples from multiple TSC patients, we found that BCs express progenitor proteins while selectively lacking markers of differentiated progeny and display heightened mTORC1-dependent activity. We also observed sample-to-sample variation in enrichment for several previously proposed BC markers, suggesting that the reported mixed-lineage phenotype of these cells is variable across patients. This study illustrates the utility of high-dimensional, protein-based imaging in better understanding possible and consistent features of diagnostic cells in human tissues.

## Results

### Balloon Identifier (BAIDEN) can be trained to identify balloon cells in tubers

As an initial-use case for the semi-automated identification of unusual cells based on histopathological criteria, we tested the ability of a trainable, machine-learning pipeline to identify the enlarged balloon cells (BCs) frequently found in brain tissue from patients with TSC. Traditionally, BCs are identified in resected epileptic foci from patients using hematoxylin and eosin (H&E) staining, which is not yet readily compatible with high-dimensional approaches such as imaging mass cytometry (IMC). BCs are often present at extremely low abundance within a given tissue, complicating efforts to select focused regions of interest for high-dimensional studies. Toluidine blue O (TBO), which has a high affinity for acidic tissue components, including nucleic acids, can be combined with IMC (Sridharan and Shankar, 2012; Baars et al., 2021). Therefore, a <u>Ba</u>lloon <u>Iden</u>tifier (BAIDEN) workflow was established to automate the identification of prospective BCs, to identify regions with high densities of such cells, and to scale these processes across the large areas included in whole-slide images (**Figure 1A**; https://github.com/ihrie-lab/BAIDEN). In brief, whole-slide images of tuber samples obtained during resections of epileptic foci were stained with TBO (compatible with IMC) and, subsequently, H&E (necessary for manual annotation), were registered and tiled. Completion of the BAIDEN workflow generates reconstructed whole-slide images with the following 4 channels: H&E, TBO alone, a grayscale probability mask identifying possible BCs, and a TBO image on which cells exceeding a user-defined probability threshold are identified and outlined. The workflow was trained using an initial set of 40 tiles from whole-slide images of tubers in which an expert neuropathologist (B.C.M.) manually identified BCs (see Materials & Methods). These training images were curated to capture various BC morphologies and some typical experimental artifacts (e.g., folded tissue). The trained workflow was then applied to a testing dataset of 432 additional annotated tiles. In this dataset, BAIDEN achieved a precision of 0.9273 and a recall of 0.6892 versus neuropathologist annotation, indicating that, with the user-defined settings specified, the workflow was highly accurate in outlining 92.73% of “true” BCs but potentially excluded some BCs (31%) that could not be assigned with high confidence (**Figure S1**). The available implementation of BAIDEN allows users to set different thresholds for labeling probable BCs on analyzed images, meaning this parameter could be adjusted if the user preference was to identify all possible BCs while potentially including some false positives (i.e., lowering precision). In this instance, rigorous identification of *bona fide* BCs was prioritized. In tandem with these analyses, tuber samples were stained and analyzed by mass cytometry. A representative, pseudo-colored image illustrates multiple antigens (7 of 36 channels, **Figure 1B**).

**Figure 1:**
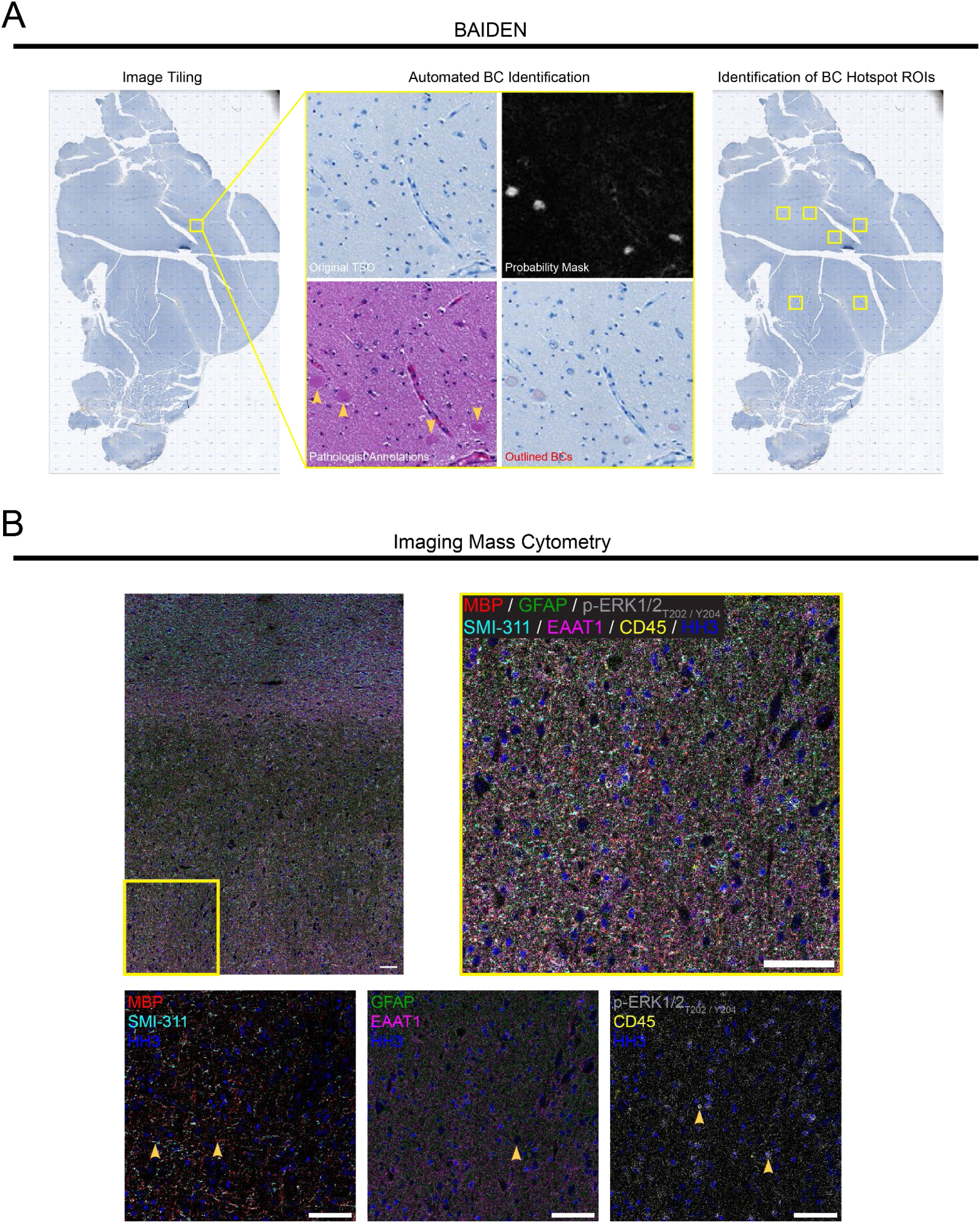
A machine-learning workflow and imaging mass cytometry to study balloon cells in tubers. A) Schematic of BAIDEN. **<u>Ba</u>**lloon **<u>Iden</u>**tifier (BAIDEN) partitions each input whole-slide image (WSI) into several tiles, each undergoing a model evaluation to probabilistically estimate the occurrence of BCs. BC hotspot regions can subsequently be identified by the user and situated in the context of the WSI for ensuing IMC analysis. B) Pseudo-colored imaging mass cytometry (IMC) image of the full imaged region of interest (ROI) for an example tuber specimen (VUT034). Yellow box delineates area detailed on the right and below. Scale bar = 100 μm for all images. 6 of 36 antigens are shown. Samples include histone H3 (HH3; blue) as the nuclear marker. Pseudo-colored images of detailed area. Yellow arrowheads denote listed antigens. Image on the left highlights MBP (red) and SMI-311 (cyan). Image in the middle highlights GFAP (green) and EAAT1 (magenta). Image on the right highlights p-ERK1/2_T202/Y204_ (grey) and CD45 (yellow).

To systematically explore previously proposed BC markers and lineage features, a customized 36-antibody panel was developed in tandem with a microarray of 21 positive and negative control samples (**Tables 1 and 2**, **Figure 2**). The antibody panel included markers for major immature and mature cell types expected to be present in normal brain (GFAP, EAAT1, SMI-311, β-III-tubulin, OLIG2, MBP, CD45, and CD31), proposed BC markers (EGFR, TTF-1 [NKX2.1], CD133, CD29 [integrin β_1_], vimentin, FAPB7 [BLBP], and EMX1), and phosphorylated proteins indicative of elevated mTOR kinase activity (p-S6_S240/S244_, p4EBP1_T37/T46_, and p-STAT3_S727_) and overlapping signaling pathways (MAPK, RSK, and EGFR). For each antigen of interest, the range and specificity of signal intensity across all control samples were used to confirm successful staining (**Figure 2, Figure S2**). Based on these initial quality-control assessments, three antigens (CD29 [integrin β_1_], PTPRZ1, and p-STAT5_Y694_) were removed from further analyses, as they did not reliably exhibit staining in positive controls or other samples.

**Figure 2:**
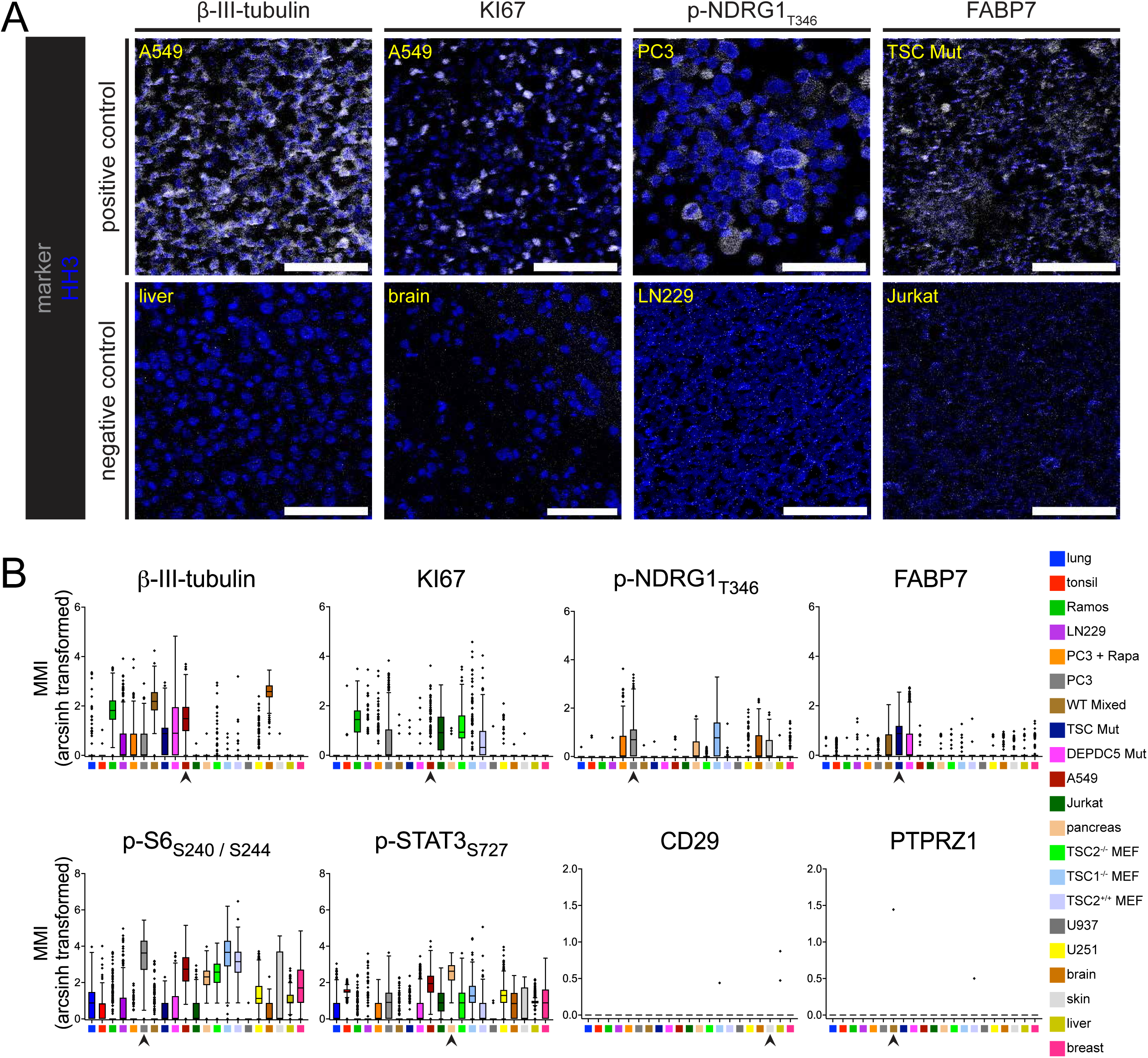
Construction and validation of custom imaging mass cytometry panel. A) Example images for individual antigens on positive and negative control cores. Scale bar = 100 μm for all images. B) Summary Tukey box-and-whisker plots of mean mass intensity (MMI) values for all events in each displayed antigen (top). Black arrowheads denote the positive core for a given antigen. Summary Tukey box-and-whisker plots of MMI values for antigens with high- and low-dynamic ranges (bottom). The legend to the far right shows the corresponding color for the control cores used.

### Balloon cells in tubers are cytomegalic and variably abundant

A set of 7 resected tuber samples was analyzed using the IMC panel detailed above. Prior to ablation for mass-based imaging, each sample was stained with TBO to enable prospective identification of BCs, and an adjacent serial section was stained with H&E (**Figure 3**) to aid in manual, retrospective annotation. Candidate BC-containing areas were first identified by running BAIDEN on whole-slide images. Ablated areas of potential interest were subsequently annotated for confirmed BCs by an expert neuropathologist (B.C.M.) using aligned TBO, H&E, histone H3 (HH3), and p-S6_S240/S244_ mass cytometry channels (**Figure 3A**). Images with expert-annotated BCs were subsequently processed to analyze all mass-tagged antibody channels using the *steinbock* pipeline (Windhager et al., 2023). Across the 7 tissue samples, 197 BCs were identified out of 16,133 cells (1.22%). The highest number of BCs identified in a tissue was 60 (DUH01; 4.86%) while the lowest number was 2 (VUT021; 0.14%). 2 of the 7 tissues (VUT021 and VUT022) were excluded from downstream analyses as they contained fewer than 5 identified BCs in the imaged region of interest. A total of 192 BCs out of 12,725 cells (1.51%) were used for downstream analyses (**Figure S5**). To support the goal of identifying antigens selectively enriched in BCs versus all other cells present in the tissue, signal intensity for each antigen, as well as a subset of scored cytoarchitectural features, was compared between BCs and non-BCs (**Figures 3 and 4; Figures S4 and S5**). As expected, BCs tended to have a larger area (**Figure S5**) and decreased eccentricity (**Figure S5**) relative to other cells within tuber samples. While BCs in some samples were enriched for specific features, a relatively limited number of channels showed consistent increases in BCs versus matched non-BCs when compared across all analyzed samples (**Figures S4 and S5**).

**Figure 3:**
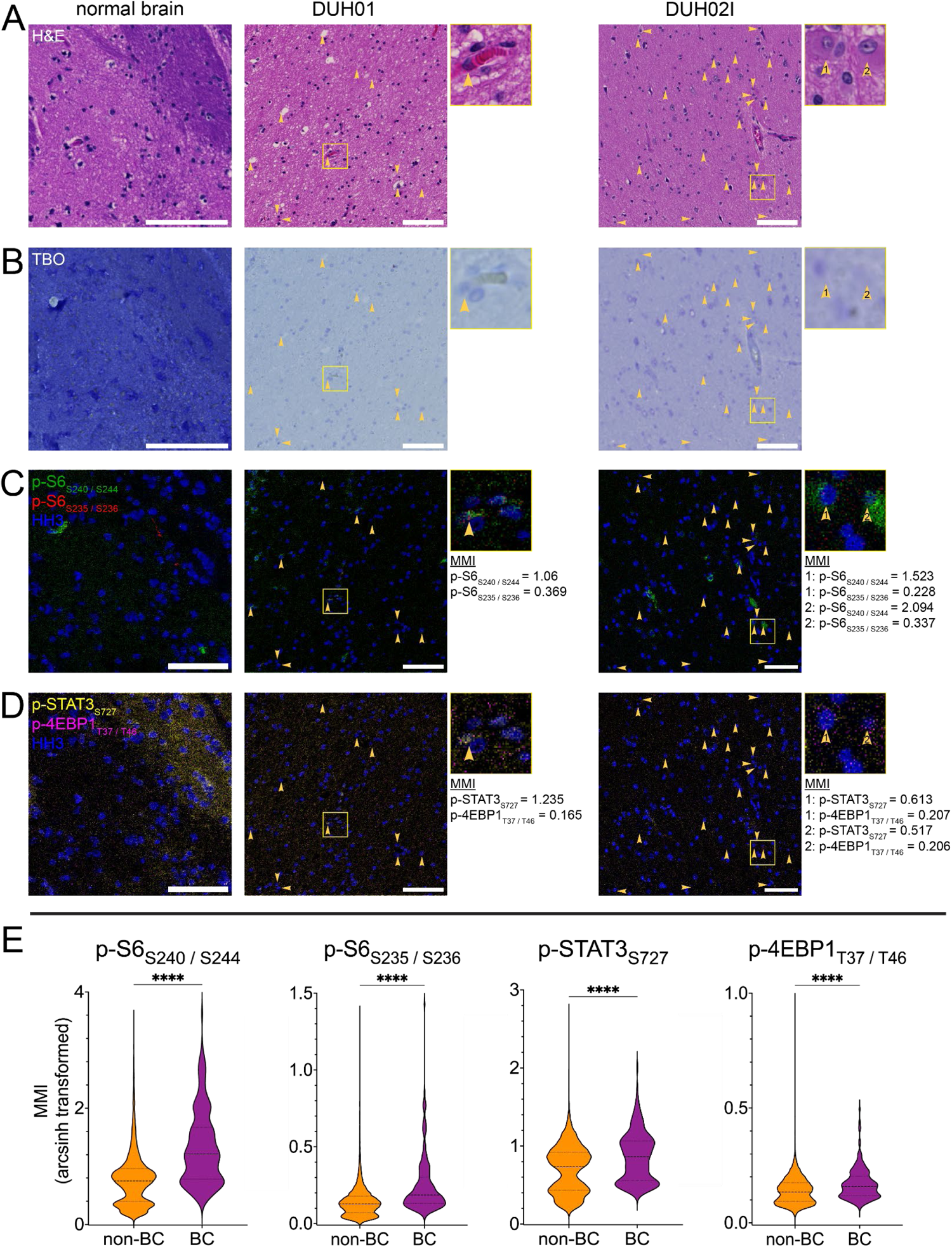
Balloon cells show elevated mTOR-dependent phosphorylation events. Rows show different stains and / or pseudo-colored phosphorylation events while the columns show normal brain and two tuber samples (DUH01 and DUH02I), respectively. A) Hematoxylin and eosin (H&E) staining of adjacent serial sections. Scale bar = 100 μm for all images. B) Toluidine blue O (TBO) staining of regions prior to ablation for IMC. C) Pseudo-colored IMC images of p-S6_S240 /S244_ (green) and p-S6_S235/S236_ (red) with HH3 (blue) as the nuclear marker. Yellow boxes are the areas of the insets to the right for all images. Yellow arrowheads denote identified balloon cells (BCs). MMI values are provided for each identified BC in the insets. D) Pseudo-colored IMC images of p-STAT3_S727_ (green) and p-4EBP1_T37/T46_ (red) with HH3 (blue) as the nuclear marker. E) Violin plots comparing hyperbolic arcsine (arcsinh)-transformed MMI values between non-BCs and BCs for p-S6_S240/S244_, p-S6_S235/S236_, p-STAT3_S727_, and p-4EBP1_T37/T46_, respectively. Heavy and light dashed lines demarcate median, 25^th^, and 75^th^ percentiles, respectively. ****p < 0.0001; two-tailed, Mann-Whitney test.

**Figure 4:**
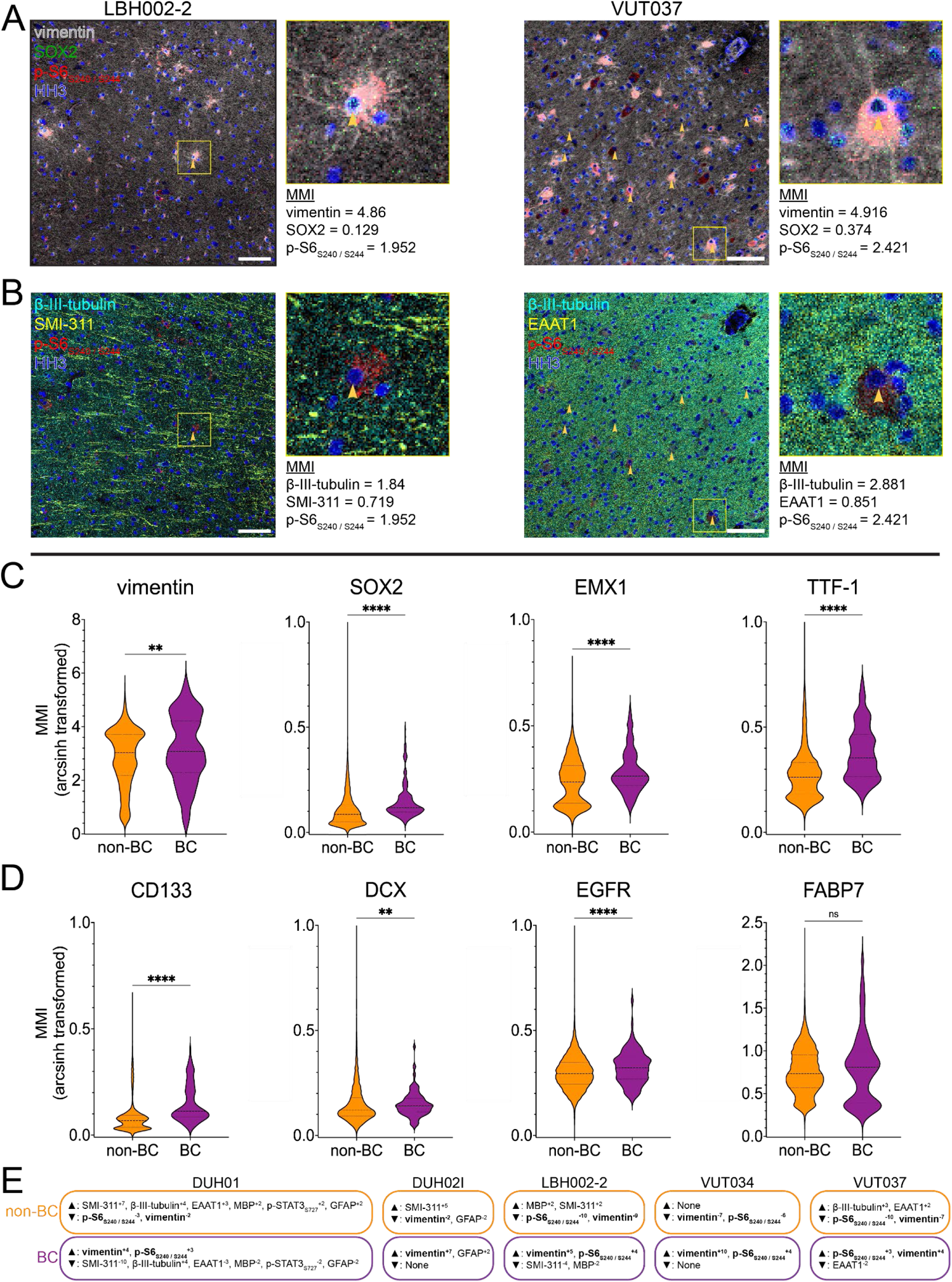
Balloon cells express progenitor-associated proteins. Rows show different proteins and phosphorylation events while the columns show two tuber samples (LBH002-2 and VUT037), respectively. A) Pseudo-colored IMC images of vimentin (grey), SOX2 (green), and p-S6_S240/S244_ (red) with HH3 as the nuclear marker. Yellow boxes are the areas of the insets to the right for all images. Yellow arrowheads denote identified balloon cells (BCs). MMI values are provided for each identified BC in the insets. Scale bar = 100 μm for all images. B) Pseudo-colored IMC images on the left depict β-III-tubulin (cyan), SMI-311 (yellow), and p-S6_S240/S244_ (red) with HH3 as the nuclear marker. Pseudo-colored IMC images on the right depict β-III-tubulin (cyan), EAAT1 (yellow), and p-S6_S240/S244_ (red) with HH3 as the nuclear marker. C) Violin plots comparing arcsinh-transformed MMI values between non-BCs and BCs for vimentin, SOX2, EMX1, and TTF-1, respectively. Heavy and light dashed lines demarcate median, 25^th^, and 75^th^ percentiles, respectively. **p < 0.01, ****p < 0.0001; two-tailed Mann-Whitney test. D) Violin plots comparing arcsinh-transformed MMI values between non-BCs and BCs for CD133, DCX, EGFR, and FABP7, respectively. Heavy and light dashed lines demarcate median, 25^th^, and 75^th^ percentiles, respectively. **p < 0.01, ****p < 0.0001, ns = not significant; two-tailed Mann-Whitney test. E) Marker enrichment modeling (MEM) labels for non-BCs and BCs for each analyzed sample. The superscripts are on a scale between −10 (selectively de-enriched; ▾) and 10 (selectively enriched; ▴). Emboldened labels highlight common labels amongst all analyzed samples.

### Balloon cells show elevated mTOR-dependent phosphorylation events

Consistent with previous reports, BCs exhibited higher per-cell levels of protein phosphorylation events associated with mTOR kinase activity. Sample-to-sample variability was observed for specific phospho-proteins, with BCs in 3 out of 5 tissues having statistically significant elevated levels of mTOR complex 1 (mTORC1) effectors p-S6_S240/S244_ (**Figure S4**), p-S6_S235/S236_ (**Figure S4**), p-4EBP1_T37/T46_ (**Figure S4**), and p-STAT3_S727_ (**Figure S4**). BCs in 3 of 5 samples showed increased levels of p-STAT3_Y705_ (**Figure S4**). There was no reproducible change in other readouts, including p-AKT_S473_ (**Figure S4 and S5**) and p-NDRG1_T346_ (**Figure S4 and S5**), which are more closely associated with mTOR complex 2 (mTORC2) activation (Schreiber et al., 2015; Szwed et al., 2021). BCs in 3 of 5 tissues had increased levels of p-ERK1/2_T202/Y204_ (**Figure S4**) while BCs in 1 tissue had decreased levels of p-ERK1/2_T202/Y204_ (**Figure S4**). Similarly, we did not reliably observe differences in per-cell levels of TFEB (**Figure S4**), which changes localization upon phosphorylation by mTOR (Alesi et al., 2021; Alesi et al., 2024). To complement per-sample comparisons, we pooled all BCs and non-BCs from the 5 studied samples and tested for significant differences in staining intensity for the proteins of interest. When aggregated across samples, BCs showed increased levels of p-S6_S240/S244_ (**Figure 3E**), p-S6_S235/S236_ (**Figure 3E**), p-4EBP1_T37/T46_ (**Figure 3E**), and p-STAT3_S727_ (**Figure 3E**).

### Balloon cells variably express progenitor-associated proteins

Multiple proteins and posttranslational protein modifications have been proposed to selectively identify BCs, ranging from those associated more strongly with progenitor lineages to those used to distinguish mature-cell types within the adult brain (Hilbig et al., 1999; Mizuguchi et al., 2002; Urbach et al., 2002; Lamparello et al., 2007; Orlova et al., 2010; Yasin et al., 2010; Ruppe et al., 2014; Li et al., 2015; Rossini et al., 2017; Day et al., 2020; Gelot and Represa, 2020; Wu et al., 2022). BCs across all tissues had decreased expression of β-III-tubulin at both the per-sample level and when aggregated (**Figures S4 and S5**), and 3 of 5 samples showed decreased expression of SMI-311 when compared on a per-sample basis (**Figures S4 and S5**). BCs, when compared on a per-sample basis, tended to show decreased levels of astrocytic markers GFAP and EAAT1 but did not show significant differences in the aggregate (**Figures S4 and S5**). 2 of 5 samples showed decreased MBP in BCs, though this difference was not significant when examined in the aggregate (**Figures S4 and S5**). BCs showed increased expression of vimentin across 4 of 5 samples (**Figure S4**) and in the aggregate (**Figure 4C**) as did CD133 (**Figure 4D**). SOX2, a transcription factor expressed in progenitor- and mature-cell types, was enriched in all per-sample comparisons (**Figure S4**) and in the aggregate (**Figure 4C**). In contrast, the progenitor marker FABP7 was enriched in 2 of 5 per-sample comparisons (**Figure S4**) but not in the aggregate (**Figure 4D**). BCs also showed slightly increased expression of EGFR (**Figure 4D**), TTF-1 (**Figure 4C**), DCX (**Figure 4D**), and EMX1 (**Figure 4C**).

### A statistical description of BC-enriched features

To identify potential BC identifiers that were conserved across cells in all 5 samples, the marker enrichment modeling (MEM) statistic was used to identify parameters that were selectively enriched or selectively de-enriched when BCs were compared to the remainder of the tissue (Diggins et al., 2017). Of the parameters measured, vimentin and p-S6_S240/S244_ were reliably enriched (positive MEM label and exponent) in BCs versus matched non-BCs across samples, (**Figure 4E**). Consistent with single-channel comparisons above, some samples also displayed enrichment or de-enrichment for additional proteins (**Figure 4E**).

## Discussion

There is widespread interest in working with patient samples to identify therapeutic targets, especially for rare diseases with limited treatment options such as tuberous sclerosis complex (TSC). In TSC, identification of balloon cells (BCs) is pathognomonic, yet BCs can vary widely in abundance both within and across surgically resected tissue sections and patients. Currently, the “gold standard” for identifying BCs relies on expert annotation by a neuropathologist using cytoarchitectural features (Urbach et al., 2002; Becker et al., 2006; Blumcke et al., 2011; Blumcke et al., 2017; Blumcke et al., 2021b). However, this process can be time-consuming and may introduce inter-scorer variability. A further challenge is the comparison of abnormal cells found in organoid or mouse models of TSC to *bona fide* BCs, including the ability to distinguish plausible BCs from the dysmorphic neurons that can also be found in tissues of patients with TSC and related disorders, emphasizing an urgent need to increase the efficiency and rapidity of identifying BCs.

We developed a trainable, machine-learning approach to efficiently identify BCs from whole-slide images stained with a colorimetric dye. Though the patient samples used in this study were cortical tissues from TSC patients, BAIDEN can be trained and tested on any tissue type or disorder, given sufficient depth in the training data to distinguish the cells of interest. One critical feature of its use is that the training, testing, and analysis datasets must be as uniform as possible in the staining protocol used for toluidine blue O (TBO) or other equivalent stains. While probable BCs could still be highlighted in the experimental samples described here, the probability threshold (i.e., the level of certainty when identifying probable BCs) was lower when staining or image quality was affected by experimental variation. In future, it will be of interest to determine if this approach can be extended to reliably identify cells of interest for applications such as laser-capture microdissection, which has been used to address the question of whether *TSC1/2* loss of heterozygosity (LOH) occurs in BCs and / or other cells found within epileptic foci (Ichikawa et al., 2005; Giannikou et al., 2016; Bongaarts et al., 2017; D’Gama et al., 2017; Blair et al., 2018; Baldassari et al., 2019; Eichmuller et al., 2022).

Histopathologically, BCs are characterized by an enlarged, ovoid soma and eccentrically placed nuclei (Urbach et al., 2002; Lamparello et al., 2007; Saxton and Sabatini, 2017; Najm et al., 2022). BCs in the patient tissues analyzed by IMC showed an increased area and were rounder than non-BCs, quantitatively supporting these metrics (Urbach et al., 2002; Becker et al., 2006; Blumcke et al., 2011; Blumcke et al., 2017; Blumcke et al., 2021b). Consistent with previous studies and case reports, BCs in the patient samples had heightened mTORC1-dependent signaling (Hilbig et al., 1999; Mizuguchi et al., 2002; Urbach et al., 2002; Lamparello et al., 2007; Orlova et al., 2010; Yasin et al., 2010; Ruppe et al., 2014; Li et al., 2015; Rossini et al., 2017; Day et al., 2020; Gelot and Represa, 2020; Wu et al., 2022). Both residues of p-S6 – p-S6_S240/S244_ and p-S6_S235/S236_ – were elevated in some BCs. The S240 / S244 residue is phosphorylated by mTOR kinase while ribosome S6 kinase (RSK) can also phosphorylate the S235 / S236 residue (Ruvinsky et al., 2005; Ruvinsky and Meyuhas, 2006; Crino, 2013). In addition, both residues of p-STAT3 – p-STAT3_S727_ and p-STAT3_Y705_ – were elevated in some BCs. While the S727 residue is primarily phosphorylated by mTOR, the Y705 residue is also phosphorylated by several other kinases and is necessary for the dimerization and transcriptional activity of STAT3 (Huang et al., 2014). The mean mass intensity (MMI) value differences between BCs and non-BCs were higher and more statistically robust for p-S6_S240/S244_ and p-STAT3_S727_, suggesting there is a preference towards mTORC1-dependent phosphorylation, consistent with the slightly elevated levels of p-4EBP1_T37/T46_.

As the understanding of BCs as a hallmark of TSC-associated FCD has grown, interest in identifying their cell of origin has increased. Prior work has postulated BCs express various proteins that distinguish progenitor cells and differentiated progeny, yet these studies largely did not assess these proteins simultaneously nor across multiple samples and patients (Hilbig et al., 1999; Mizuguchi et al., 2002; Urbach et al., 2002; Lamparello et al., 2007; Orlova et al., 2010; Yasin et al., 2010; Ruppe et al., 2014; Li et al., 2015; Rossini et al., 2017; Day et al., 2020; Gelot and Represa, 2020; Wu et al., 2022). BCs in the patient tissues examined here tended to express proteins associated with progenitor-cell identity and function rather than mature-cell identity. Specifically, vimentin, SOX2, CD133, and EGFR were elevated in BCs while markers used to identify mature cells, including β-III-tubulin, SMI-311, GFAP, and EAAT1, tended to be decreased. In the developing brain, regional patterning is associated with the generation of specific neuronal subtypes and can be detected via staining for region-specific transcription factors at specific developmental stages. TTF-1, which distinguishes the medial ganglionic eminences (ventral structures), has previously been proposed as a marker largely absent from tubers (Rushing et al., 2019). Interestingly, the BCs analyzed by IMC had slightly increased expression of both TTF-1 and the transcription factor EMX1 (typically prevalent in cortex), though this varied across samples. BCs also had increased expression of DCX, which distinguishes migratory immature neurons (Gleeson et al., 1999; Ayanlaja et al., 2017). Collectively, these patterns of expression, as well as the MEM scores for BCs in each sample, suggest that while BCs reflect an immature and / or mixed-lineage phenotype, relatively few proposed markers of this cell type are uniformly expressed from cell to cell or patient to patient. Identification of uniform and variable features may help in determining which signaling arms of this pathway can and should be targeted in tubers and BCs – a relevant question as not all mTOR inhibitors in development for the clinic have equivalent efficacy on downstream phosphorylation events (Choi et al., 1996; Fan et al., 2018; Mao et al., 2022). As in studies of other progenitor populations, examining multiple parameters will likely be critical to confirming BC, or BC-like, identity in cell-based models.

Some challenges and important considerations were evident in these initial studies. First, despite the uniform preparation and staining of all tissue sections and controls with a single antibody cocktail, we did observe sample-to-sample variation in the staining intensity, as evident in per-sample comparisons. This variation could be due to variations in sample fixation and processing and may also reflect regional variation within cortical tissue, as tubers can present in many locations in the brain and display variability in their radiographic and pathologic features (Jesmanas et al., 2018). Nonetheless, systematic comparisons across tissue samples were able to highlight markers with uniform direction of variation across patients (i.e., proteins that were always enriched, or always de-enriched, in BCs versus non-BCs). These comparisons could be further refined by systematically identifying suitable normal cell comparators within the tissue of interest. Second, carefully generating standardized controls for custom antibodies was valuable in identifying the dynamic range of signal for each target and confirming the lack of signal in cells of interest (as in the case of mature-cell markers in BCs). Collectively, these data present a panel development, validation, and analysis strategy that can be applied to the brain and many other tissues where high-dimensional imaging is of interest.

## Contributions

JSA: sample collection, data acquisition, data analysis & interpretation, writing – original drafts

AAB: sample collection, data acquisition, data analysis & interpretation, writing – original drafts

RK: data acquisition, data analysis & interpretation, writing – original drafts

LCG: data acquisition, experimental design

MBLC: sample collection, data acquisition, experimental design

MV: supervision, sample collection

MZ: supervision, sample collection

SW: supervision, sample collection

BCM: data acquisition, data analysis & interpretation

KCE: supervision, conceptualization, funding acquisition, sample collection, writing – final version

RAI: supervision, conceptualization, funding acquisition, data analysis & interpretation, writing – original drafts & final version

All authors: review & approval of manuscript

## Acknowledgements

We thank patients and their families for agreeing to participate in tissue collection for research. Research was supported by a pilot award from the Vanderbilt Brain Institute, the National Institutes of Health R01NS118580 (RAI and KCE) and a supplement to U54CA217450 (RAI), T32 GM07628 and F31 NS120608 (LCG), T32 HD007502 and R01NS118580-S1 (MBLC), the Ben & Catherine Ivy Foundation (RAI), the Meharry Medical College MD/PhD Endowment Grant (JSA), and a gift from the Michael David Greene Brain Cancer Fund at the Vanderbilt–Ingram Cancer Center (RAI, AAB). The VUMC Translational Pathology Shared Resource (TPSR) is supported by NCI/NIH Cancer Center Support Grant P30CA068485.

## Materials & Methods

### BAIDEN

<u>Ba</u>lloon <u>Iden</u>tifier (BAIDEN; https://github.com/ihrie-lab/BAIDEN) is a pixel-based classifier that identifies balloon cells (BCs) from input whole-slide images of tuber specimens stained with toluidine blue O (TBO). Generally, BAIDEN operates at the tile level and stitches together its outputs to form a new multi-channel, whole-slide image composed of the original TBO stain, the model’s BC probability estimates, and outlined BCs overlaid onto the TBO image that survive quality control.

The underlying model was developed using Ilastik (Berg et al., 2019), a flexible machine-learning toolkit that utilizes random forests to estimate segmentation masks based on pixel-level features related to color and intensity, edges, and texture.

After probabilistically estimating the occurrence of BCs, BAIDEN feeds the generated segmented maps into CellProfiler (Carpenter et al., 2006) for finer object detection. In a sequential fashion, the outputs from Ilastik are smoothened with a Gaussian kernel (σ = 1) and subjected to primary object identification in which red outlines are constructed around the most putative BCs with the appropriate diameter (between 15 and 100 pixels). These outlines are then overlaid onto the original TBO image to ease visualization of identified BCs for downstream regions of interest (ROI) identification for imaging mass cytometry (IMC).

### Immunostaining

#### Blocking Solution

The blocking solution for all immunostaining approaches was 1X PBS (no Ca^2+^ / Mg^2+^) with 1% normal donkey serum (Fisher Scientific; OB003001), 1% bovine serum albumin (Sigma-Aldrich; A7906), 0.1% Triton X-100 (Fisher Scientific; AAA16046AE), and 0.01% sodium azide (Fisher Scientific; BP922I) as a preservative. The blocking solution was stored at 4°C between uses.

#### Antigen Retrieval and Slide Processing

Slides were deparaffinized, rehydrated, and subjected to heat-induced epitope retrieval using Trilogy (Sigma-Aldrich; 920P-09) in a Cuisinart pressure cooker following the manufacturer’s directions. Briefly, slides were incubated at a high-pressure setting for 10 minutes, followed by manual pressure release and a 5-minute “hot rinse” in a second jar of Trilogy before rinsing with ddH_2_O. For paraffin-dipped slides, individual slides were first dewaxed in separate Coplin jars with 2 xylene washes for 30 minutes each and then underwent an additional Trilogy “hot rinse” for 5 minutes before the final “hot rinse” step.

#### Blocking and Antibody Incubation in Humid Chamber (Fluorescence Imaging)

Slide-mounted samples were outlined with a hydrophobic marker and placed horizontally in a humid chamber for all staining and washing steps. Staining followed standard methods. Briefly, samples were rinsed 3 x 5 minutes with 1X PBS before incubation with blocking solution for at least 30 minutes at room temperature. Next, primary antibodies (listed in **Table 1**) diluted in the blocking solution were added to samples and incubated overnight at 4°C. The following day, samples were rinsed 3 x 5 minutes with 1X PBS before incubation with secondary antibodies for an hour at room temperature. For initial antibody validations using fluorescent detection, Hoechst (10 mg/mL; Fisher Scientific; PI62249) was added to the secondary antibody cocktail. Samples were rinsed 3 x 5 minutes with 1X PBS before mounting with Fluoromount G (Fisher Scientific; 50-259-73).

#### Blocking and Antibody Incubation in Parhelia Chamber (Fluorescence Imaging)

Following dewaxing and antigen retrieval, slides were prepared for blocking by placing about 30 mL of ddH_2_O into the Parhelia S-Type OmniStainer (Parhelia Biosciences; 40002). Slides were immersed in ddH_2_O, and a freshly prepared covertile (Parhelia Biosciences; 40200) was applied while the slides were immersed. All subsequent solutions were applied by adding 150 μL of solution slightly (∼2 mm) above the interface between the covertile and the slide. Slides were rinsed 3 x 5 minutes with 1X PBS, followed by incubation with the blocking solution alone for at least 30 minutes at room temperature. Primary antibodies (listed in **Table 1**) diluted in the blocking solution were applied, and samples were incubated overnight at 4°C. The following day, samples were rinsed 3 x 5 minutes with 1X PBS. Secondary antibodies diluted in Hoechst-containing blocking solution were added for 1 hour at room temperature. Samples were rinsed 3 x 5 minutes with 1X PBS, removed from the Parhelia chamber, and coverslips were mounted with Fluoromount G.

#### Toluidine Blue O Incubation

Slides were prepared for toluidine blue O (TBO; Sigma-Aldrich; T3260) staining in a Parhelia OmniStainer as described above. After 3 x 5-minute rinses in 1X PBS, samples were incubated with 1% TBO (w/v) for 5 minutes at room temperature. Samples were rinsed 1 x 5 minutes with 1X PBS, removed from the Parhelia chamber, and coverslips were mounted with aqueous mounting media (1:1 1X PBS:glycerol) and used for whole-slide, brightfield imaging.

#### Blocking and Antibody Incubation in Parhelia Chamber (Metal-Tagged Antibody Staining) and Imaging Mass Cytometry

Following TBO staining and imaging, coverslips and mounting media were soaked off in warm water. An OmniStainer covertile was applied, and slides were rinsed one time each with ddH_2_O, 1X PBS, and blocking solution. Fresh blocking solution was added and incubated for 30 minutes at room temperature. Primary antibodies were applied as described above and incubated overnight at 4°C. The following day, samples were rinsed 3 x 5 minutes with 1X PBS and 1 x 5 minutes with ddH_2_O. The covertile was removed, and slides were allowed to dry overnight before imaging on a Hyperion Imaging System (Washington University in St Louis Human Immunology Core). Tissue was laser ablated at 200 Hz, and data were exported as .mcd and .txt files.

#### Metal-Tagged Antibodies

All antibodies used for IMC analysis are listed in **Table 1**. Custom-conjugated antibodies (emboldened in **Table 1**) were prepared using standard methods (Bendall et al., 2011) with Maxpar X8® Chelating Polymer Kits (Standard BioTools; 201300).

### Specimens

#### Human Specimens and Tissue Processing

Tissues were collected from Vanderbilt Children’s Hospital between 2018 and 2023 under IRB #130550, from Duke University Hospital in 2022 under IRB #108389, and from Le Bonheur Children’s Hospital between 2022 and 2023 under IRB #20-07761-XP with written informed consent in accordance with the Declaration of Helsinki. Samples were deidentified before processing. A complete list of patient demographics and clinical characteristics can be found in **Table 3**. Tubers and perituberal tissues were obtained within 1 hour of resection. Samples were processed into slabs of tissue ∼5-10 mm thick and placed in 10% neutral buffered formalin (Fisher Scientific; 426-796) for 24-36 hours at 4°C prior to transfer to 70% ethanol. Processing into paraffin, hematoxylin and eosin (H&E) staining, and serial sectioning of samples were performed by the Translational Pathology Shared Resource (TPSR) at Vanderbilt University Medical Center.

### Image Processing for Display

#### Raw Image Export

Images from the Hyperion Imaging System were exported as single-channel TIFFs using MCD Viewer (Standard BioTools) or histoCAT++ (https://github.com/BodenmillerGroup/histoCAT3D).

#### Arcsinh Transformation, Tiling, and Export

To best separate signal and noise using known positive and negative control samples (**Figure S2 and Table 2**), an in-house, Python script processed raw images and transformed the images with an inverse hyperbolic sine (arcsinh) function, using expert-selected cofactors for each metal-tagged antibody (**Table 1**). These cofactors were applied in the Python, histoCAT++, and *steinbock* analyses discussed below.

#### Image Registration

IMC images were registered with brightfield TBO images by isolating the approximate area imaged by IMC within the brightfield slide scan through consultation of an annotated thumbnail image depicting the imaged region by IMC. The resolution of the brightfield TBO image was reduced to match the resolution of the IMC image using ImageJ / FIJI (https://fiji.sc/), and the size transformation was calculated using the per-pixel scale of each image. A pixel-to-pixel registration between the TBO and IMC images was achieved using the histone H3 (HH3) IMC channel overlayed onto the downscaled, brightfield image. Rigid registration was sufficient to achieve near-pixel-perfect registration across the imaged region of interest (ROI). Adjacent H&E was manually registered to TBO by aligning tissue edges and macroscopic features in ImageJ / FIJI.

#### Image Annotation

An expert neuropathologist viewed stacked images of the H&E, TBO, HH3, and p-S6_S240/S244_ and annotated probable BCs using the ‘MultiPoint’ tool in ImageJ / FIJI.

### IMC Analysis

#### steinbock Processing and Export

Raw .mcd files were processed by *steinbock* as the Docker container according to the Bodenmiller Lab instructions (https://bodenmillergroup.github.io/steinbock/latest/) and as reported (Windhager et al., 2023). Briefly, command-line usage was used to classify pixels as either nucleus, cytoplasm, or background and produce a probability map (ilastik); segment individual cells according to the probability map produced from ilastik (CellProfiler) and produce cell masks; measure mean intensities from each identified cell according to the cell masks from CellProfiler; and export .csv files (Carpenter et al., 2006; Berg et al., 2019).

#### Spillover Correction (Data Acquisition and Application)

The spillover matrix (SM) slide (**Supplemental Table 1**) was generated and analyzed according to the Bodenmiller Lab instructions (https://bodenmillergroup.github.io/steinbock/latest/) and as previously reported (Chevrier et al., 2018). The CATALYST package in R was used to import the individual .txt files for each spotted, metal-tagged antibody, create and apply the SM, and export the SM as a .csv and stacked TIFFs for each imaged ROI (Nowicka et al., 2017).

#### Marker Enrichment Modeling (MEM) Label Processing

The marker enrichment scores of BCs and non-BCs were identified per sample using the MEM package in R, where MEM labels were generated using all proteins included in **Table 1**, and the display threshold was set at 2 (Diggins et al., 2017).

### Statistical Analysis

Statistical analysis was performed in R version 4.3.1, GraphPad Prism version 10.1.1, or Python version 3.10.4, where indicated. All analyses were graphed in R, Prism, or Python. Statistical analysis of aggregated, individual BCs and non-BCs from different patient samples was performed using a two-tailed, Mann-Whitney test. Bonferroni correction was applied, resulting in a p-value significance cutoff of 0.002 for the violin plots. Statistical analysis of paired BCs and non-BCs from different patient samples was performed using a two-tailed, Mann-Whitney test. Bonferroni correction was applied, resulting in a p-value significance cutoff of 0.0003 for summary Tukey box-and-whisker plots. Unless otherwise indicated, p-values less than 0.05 were considered significant for all statistical tests.

---

**Figure S1:**
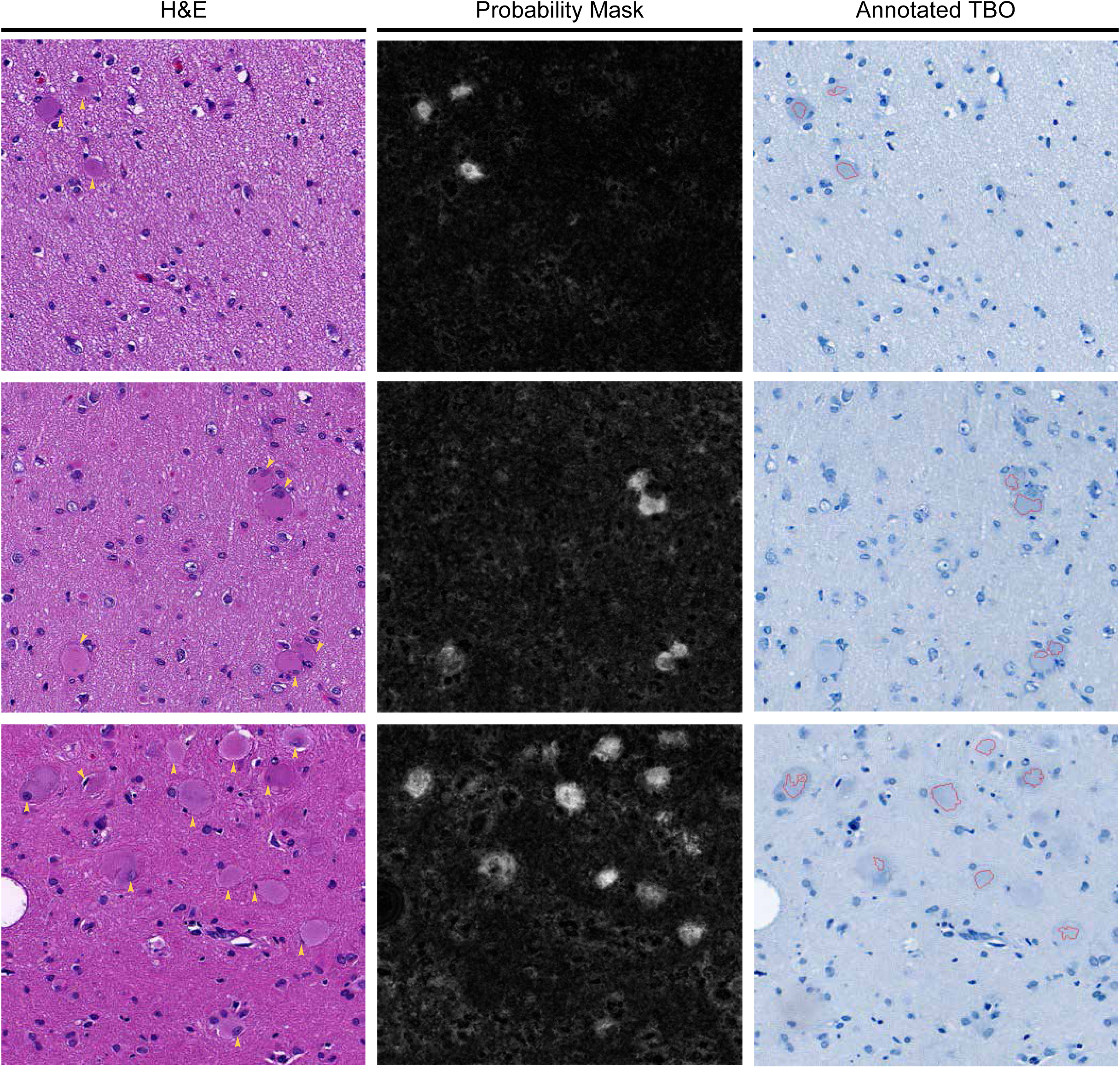
BAIDEN Performance. Comparison of neuropathologist-annotated hematoxylin and eosin (H&E) tiles with BAIDEN’s estimates. BAIDEN incurs relatively few false positives (FPs) but fails to identify every BC, resulting in a precision of 0.9273 and a recall of 0.6892.

**Figure S2:**
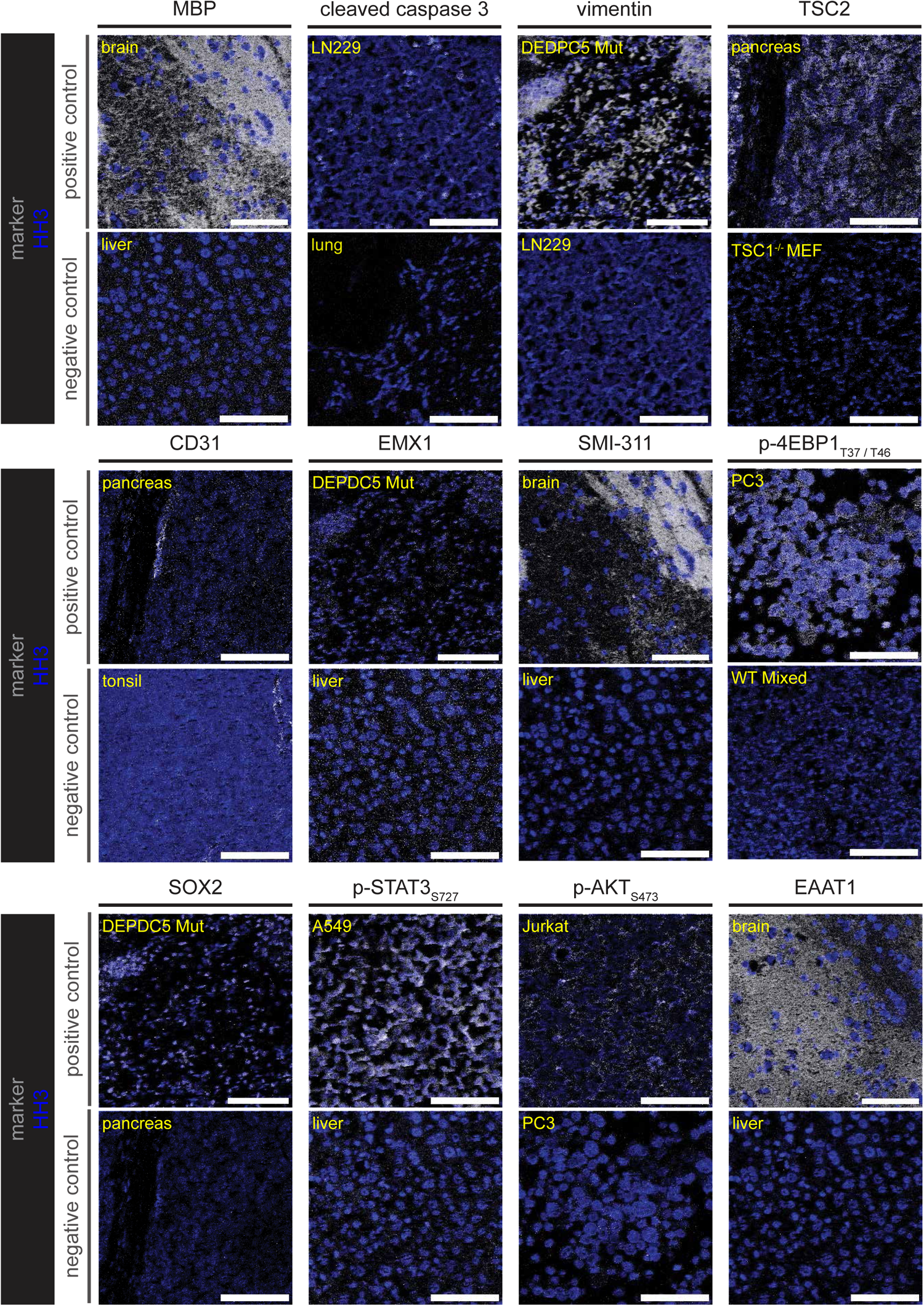
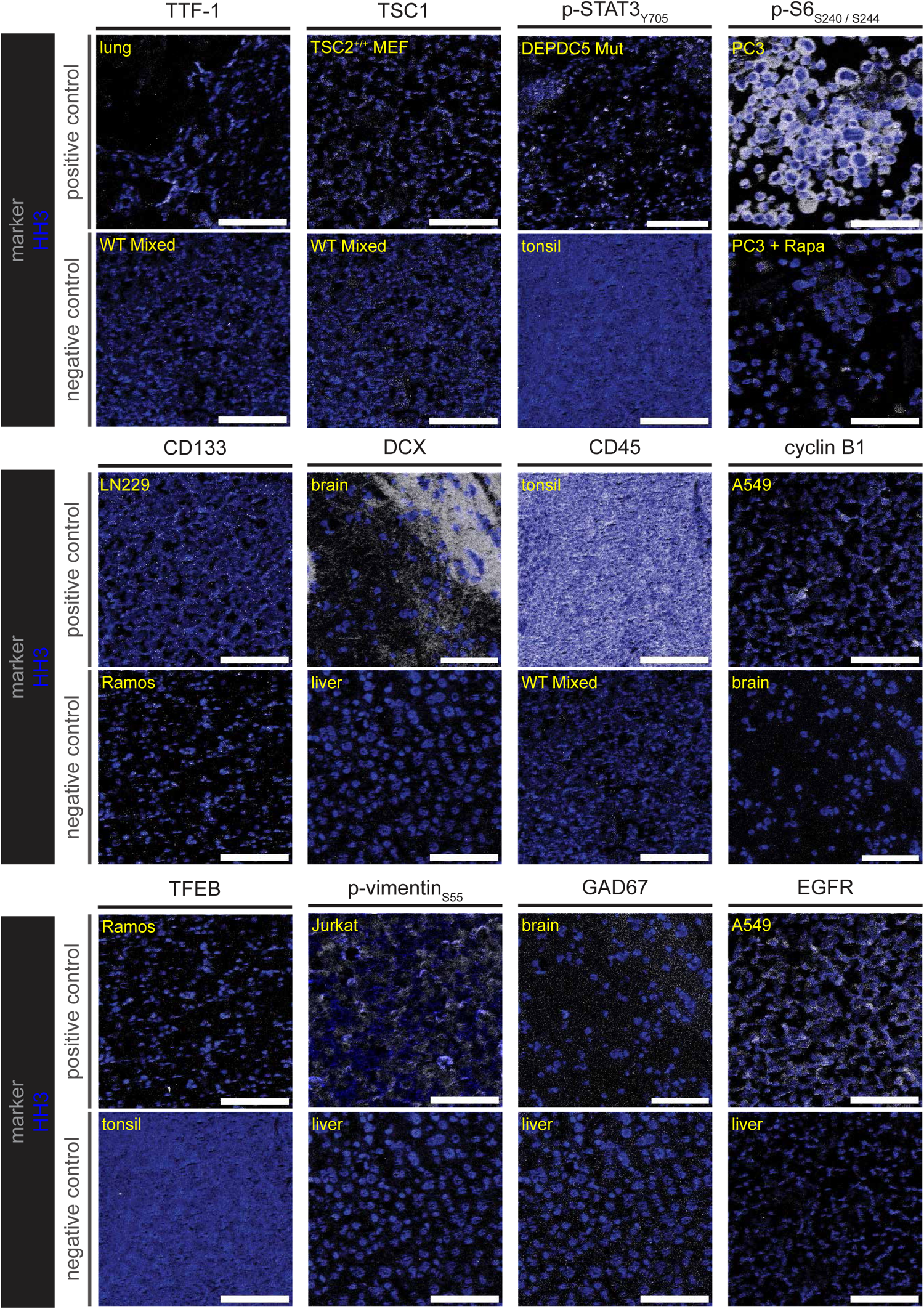
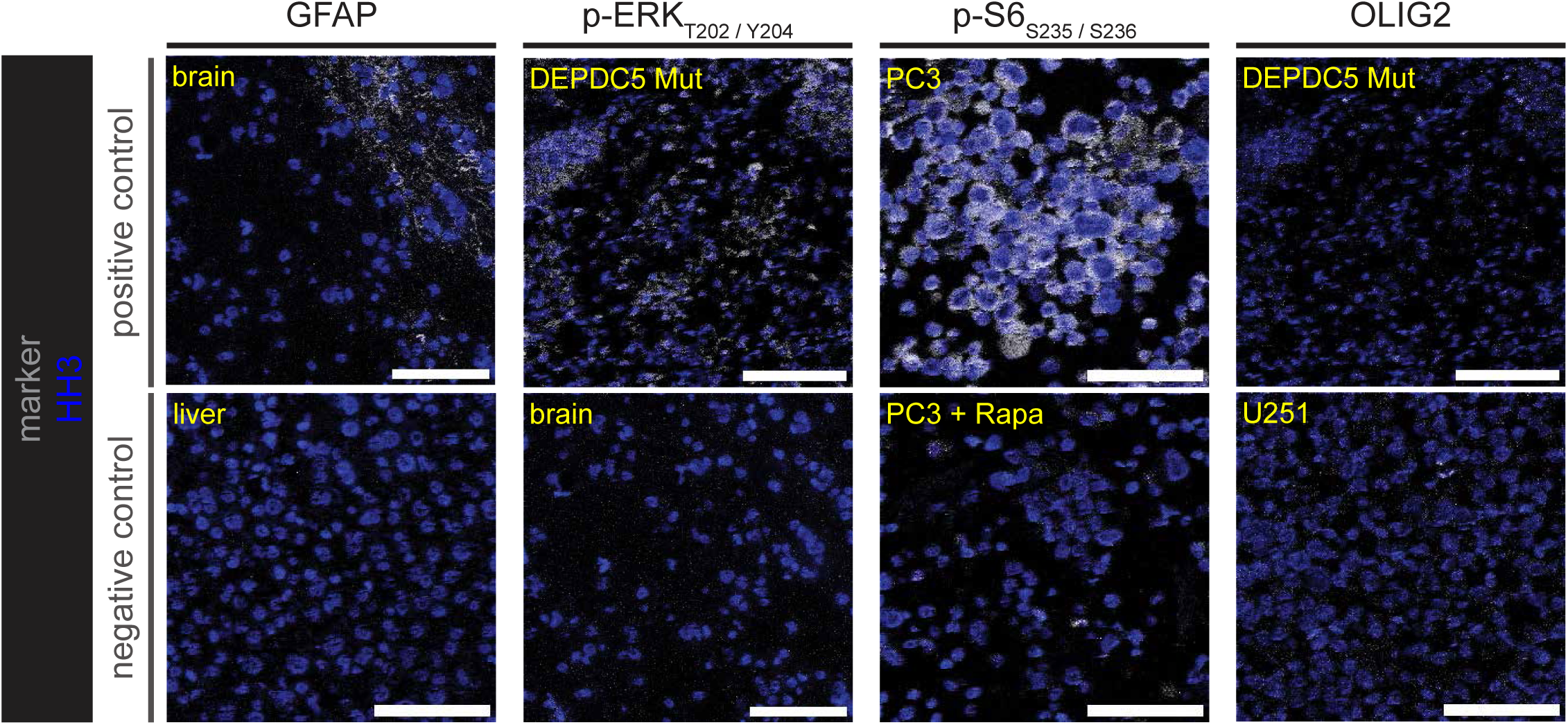
A control core microarray for validation of antibodies. Rows show a paired positive (upper) or negative (lower) control core for a given protein or phosphorylation event (arranged by columns). Protein or phosphorylation event is pseudo-colored in grey while HH3 is pseudo-colored in blue. Yellow text in the top left corner of each image indicates the control core shown. Scale bar = 100 μm for all images.

**Figure S3:**
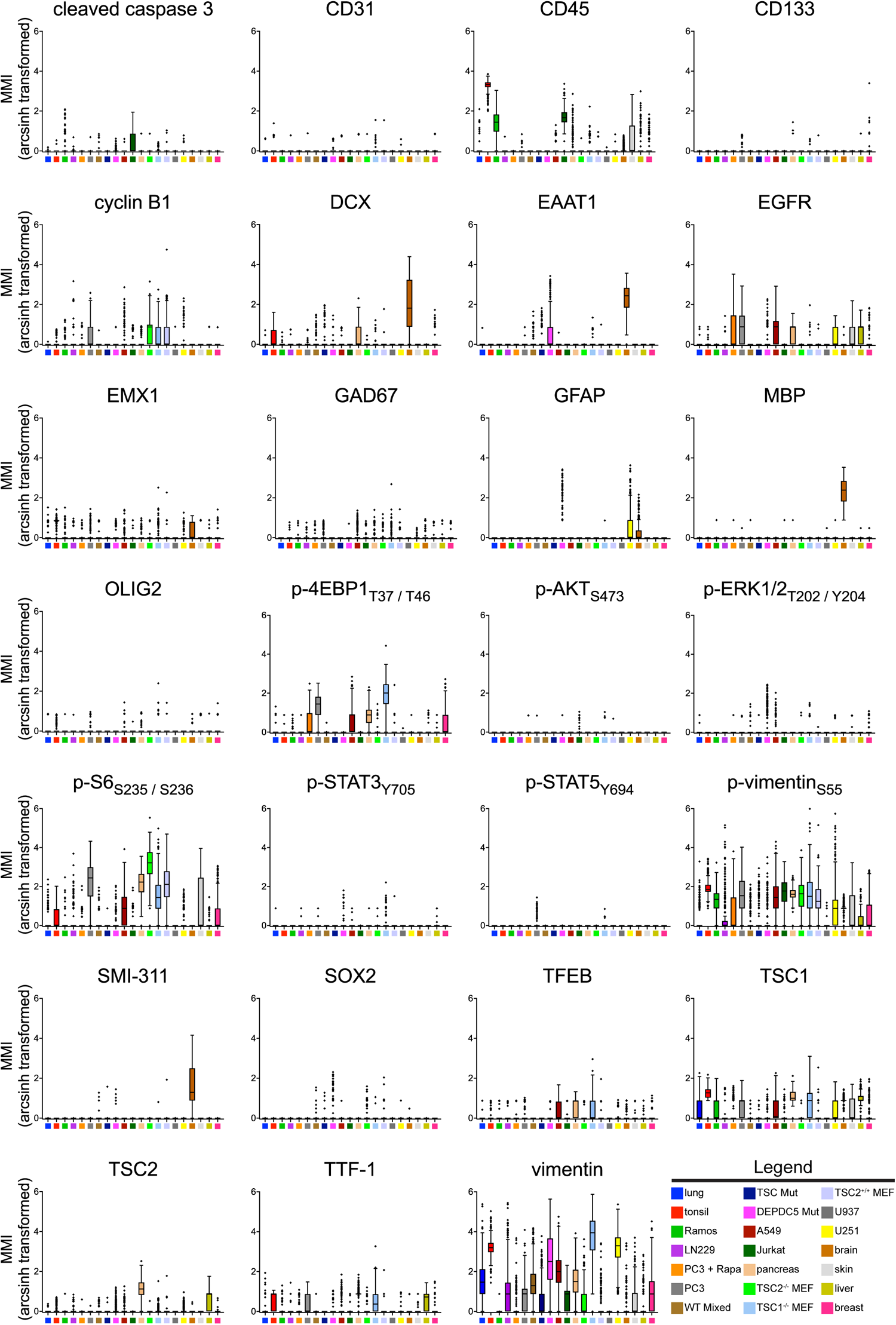
Compiled mean mass intensity (MMI) values for the control core microarray. Summary Tukey box-and-whisker plots of arcsinh-transformed MMI values of the remaining twenty-seven proteins and / or phosphorylation events for the control cores. Heavy line demarcates median while bars show maximum and minimum.

**Figure S4:**
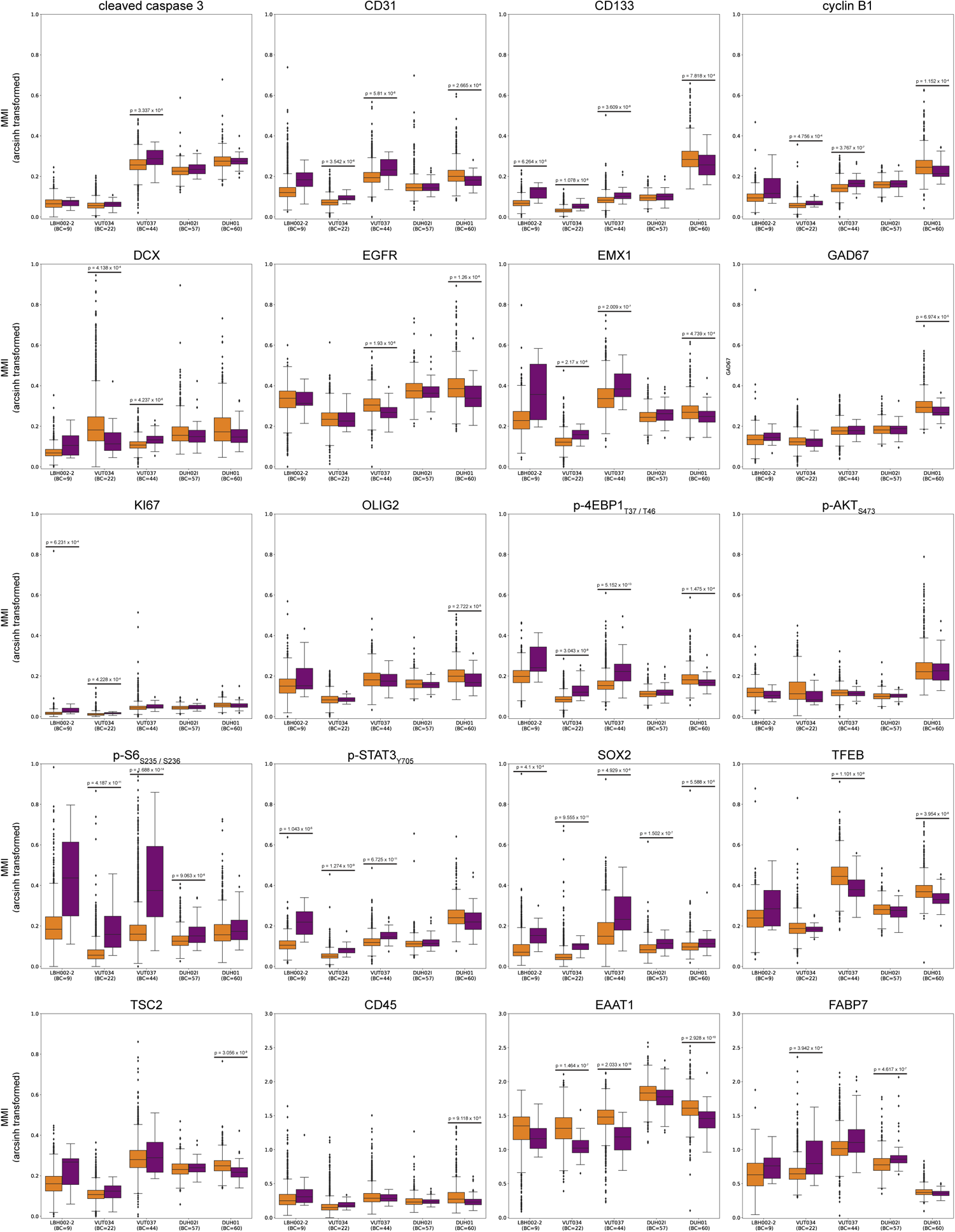
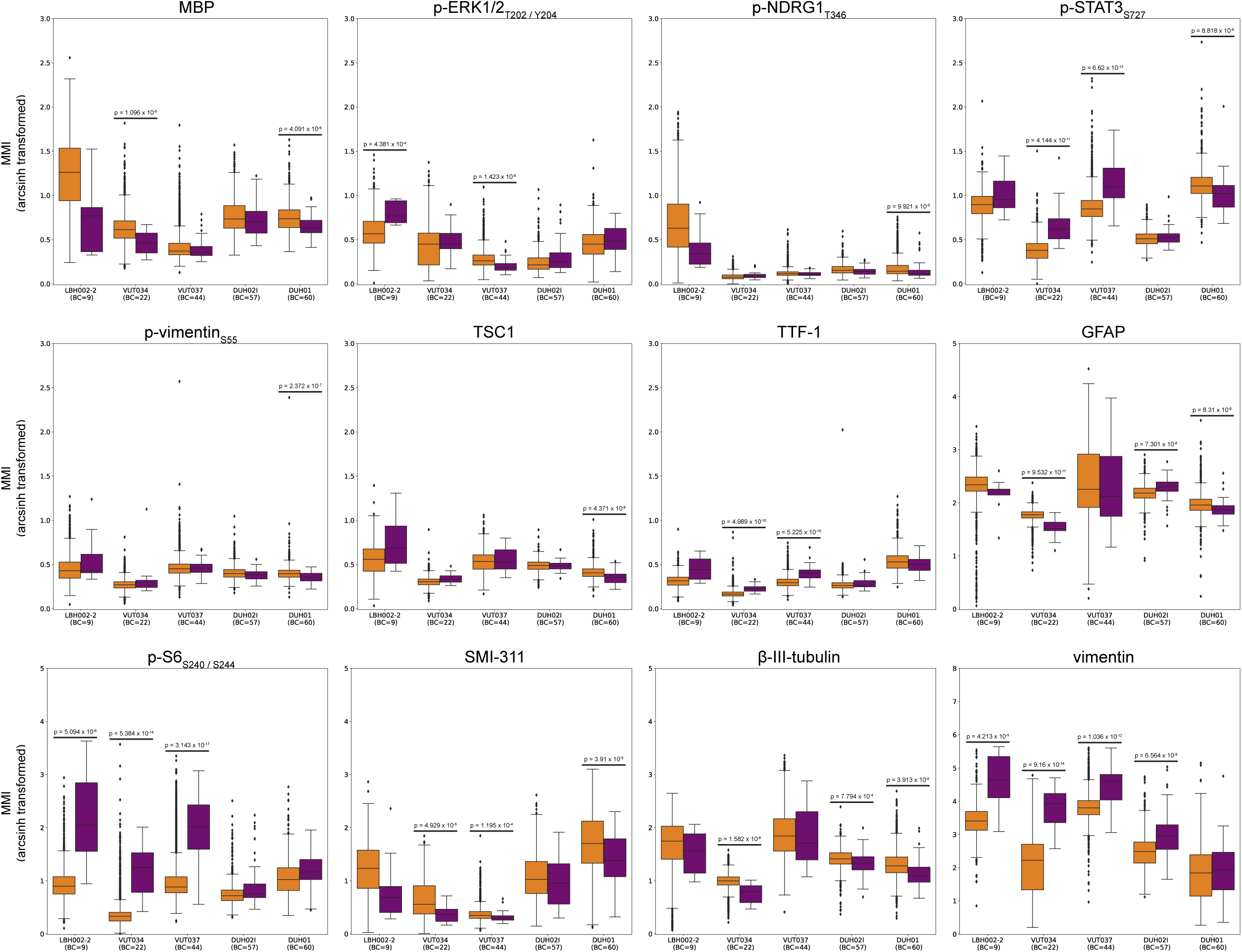
Aggregated, per-sample mean mass intensity (MMI) values for the analyzed patient samples. Summary Tukey box-and-whisker plots comparing arcsinh-transformed MMI values between non-BCs and BCs of thirty-two collected proteins and / or phosphorylation events. Heavy line demarcates median while bars show maximum and minimum. Bonferroni correction was applied, resulting in a p-value significance cutoff of 0.0003.

**Figure S5:**
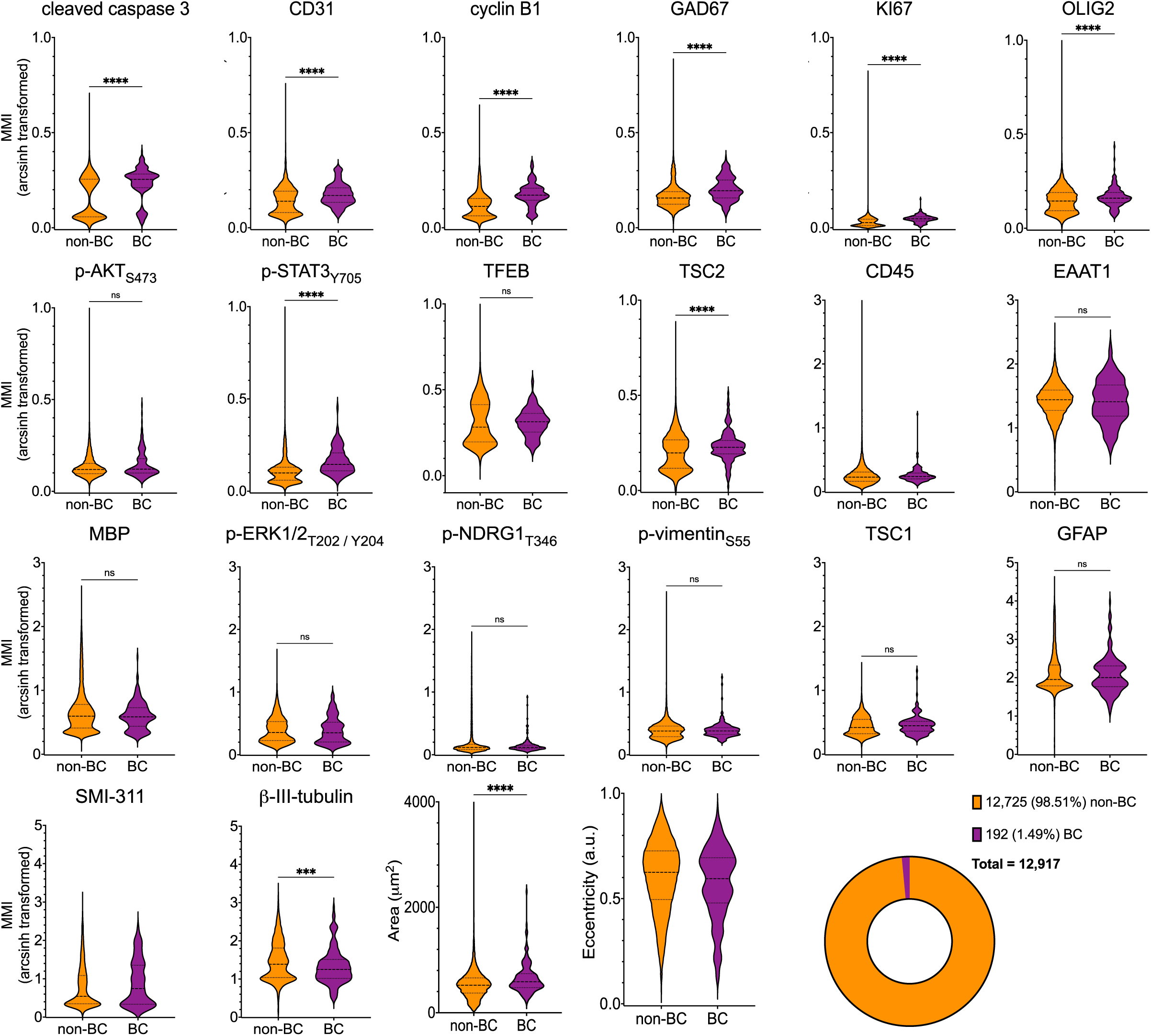
Aggregated, per-cell descriptors for the analyzed patient samples. Violin plots comparing arcsinh-transformed MMI values between non-BCs and BCs for the remaining twenty-three descriptors of the analyzed patient samples. Heavy and light dashed lines demarcate median, 25^th^, and 75^th^ percentiles, respectively. Bonferroni correction was applied, resulting in a p-value significance cutoff of 0.002. *p < 0.05, **p < 0.01, ***p < 0.001, ****p < 0.0001, ns = not significant; two-tailed Mann-Whitney test.

